# BIRC5 Regulates IGFBP-3-Induced Autocrine Loop in Cancer Stem Cells

**DOI:** 10.1101/2023.03.09.531969

**Authors:** Yeon-Jee Kahm, Uhee Jung, Rae-Kwon Kim

**Affiliations:** Department of Radiation Biology, Environmental Safety Assessment Research Division, Korea Atomic Energy Research Institute, Yuseong-Gu, Daejeon, 34057, Korea; Department of Radiation Science and Technology, Korea University of Science and Technology, Yuseong-Gu, Daejeon, 34057, Korea

**Author notes:** Corresponding author. Address: Department of Radiation Biology, Environmental Safety Assessment Research Division, Korea Atomic Energy Research Institute, 111, Daedeok-Daero 989 Beon-Gil, Yuseong-Gu, Daejeon, Korea. These authors contributed equally to this work (Yeon-Jee Kahm, Rae-Kwon Kim).

## Abstract

Baculoviral inhibitor of apoptosis repeat-containing 5 (BIRC5) is also known as survivin. BIRC5 is overexpressed in various carcinomas and is involved in cell growth and apoptosis. BIRC5, a member of the apoptosis inhibitor (IAP) family, negatively regulates apoptosis or programmed cell death by inhibiting caspase activation. Due to these properties, overexpression of BIRC5 enables specific survival and division associated with cancer malignancies. In addition, BIRC5 is highly expressed in stem cells, but not present at all in terminally differentiated cells. On this basis, there is speculation that BIRC5 may be involved in the regulation of cancer stem cells (CSCs), but few study results have been reported. In addition, the molecular mechanisms of BIRC5 regulation are not yet well understood. Through the present study, it was confirmed that BIRC5 is a key factor regulating CSCs and epithelial to mesenchymal transition (EMT) phenomena. BIRC5 was simultaneously overexpressed in lung cancer stem cells (LCSCs) and glioma stem cells (GSCs), and when the expression was suppressed, the characteristics of CSCs disappeared. In addition, IGFBP-3, a secreted factor regulated by BIRC5, is involved in signaling mechanisms that regulate cancer stem cells and EMT, and IGFBP-3 forms an autocrine chain. Based on these results, BIRC5 is proposed as a novel therapeutic target protein for LCSCs and GSCs.

## Introduction

Baculoviral inhibitor of apoptosis repeat-containing 5 (BIRC5), also known as survivin, is a member of the apoptosis inhibitor (IAP) family and negatively regulates apoptosis or programmed cell death by inhibiting caspase activation (1). It has been reported that BIRC5 has low expression in normal tissues and high expression in malignant cancer (2–4). Recently, BIRC5 was identified as an important factor in GSC, and the efficacy of materials targeting it has been reported (5). According to the aforementioned study, BIRC5 plays an important role in GSC formation, and it was reported that substances targeting it inhibit GSC formation. However, studies on BIRC5 as a regulator of CSC are not yet well known.

Cancer stem cells have been proposed as therapeutic targets by many researchers. In particular, they have been reported to be resistant to chemotherapy and radiation (6,7) and therefore are considered a major cause of difficulties in cancer treatment (8). Cancer stem cells use different marker proteins for each tissue (9). Sox2, Oct4, and Nanog are representative factors regulating cancer stem cells (10–12) and are well known as markers and regulators of stem cells (9,10). Factors that regulate stem cells are also overexpressed in cancer stem cells (13). Therefore, cancer stem cells are capable of self-renewal, which is one of the characteristics of stem cells (14). It has been reported that this self-renewal property makes cancer treatment difficult and causes cancer metastasis (15). Cancer stem cells self-renew at the metastatic site to create new cancer tissues (16). These cancer tissues often have more malignant traits than the first cancer tissues (17). In addition, cancer stem cells that survive cancer treatment regenerate on the spot and grow cancer tissue again (18). This is known as a relapse. It has been reported that cancer stem cells are involved in recurrence and metastasis.

IGFBP-3 mainly binds to IGF-I and IGF-II and is known as a major transporter of IGF. In general, IGFBP-3 is known as a protein that binds to IGF and regulates the half-life of IGF. It has also been reported that the secretion of IGFBP-3 in cancer is higher in more aggressive ER cell lines than in ER^+^ cells in breast cancer (19). In addition, it is known that the higher the concentration of IGFBP-3 in breast tissue is, the higher the risk of death (19) and the higher the expression in malignant tissue (20, 21). IGF and IGFBP are co-transported from the circulatory system to tissues, where systemic and regional regulatory systems are activated. The systemic regulatory system is the circulating IGF system and is influenced by systemic conditions such as growth hormone, insulin, and nutrition. Regional regulatory systems arise from the combination of several locally expressed components and are controlled by factors specific to each individual tissue (22).

This study investigated the critical role of BIRC5 in LCSCs and GSCs. In addition, it was identified that IGFBP-3 is regulated by BIRC-5. Therefore, BIRC5 is thought to be a potential target for new therapeutic agents.

## Materials and Methods

### Cell culture

A-549 (CCL-185^™^) and NCI-H460 (HTB-177^™^) human lung cacer cell lines were purchased from the ATCC and grown using RPMI 1640 MEDIUM(1X) (cat.no. SH30027.01; Hyclone; Cytiva). Human glioblastoma cell line, U-87 MG (30014) was obtained from the Korea Cell Line Bank and cultured in DMEM/HIGH GLUCOSE (cat. no. SH30243.01; Hyclone; Cytiva). These mediums were supplemented with 10% FBS (cat. no. SH30919.03; Hyclone; Cytiva) and 1% Penincillin-Streptomycin Solution (cat.no. SV30010; Hyclone; Cytiva). All cells were cultured at a humidified incubator with 37°C in 5% CO_2_.

### Chemicals and antibodies

All chemicals for the experiments were reagent grade or better. Antibodies against CD44 (cat. no. 3570; Cell Signaling Technology, Inc., 1:1000), Sox2 (cat. no. 3579; Cell Signaling Technology, Inc., 1:1000), Oct-4 (cat. no. 2750; Cell Signaling Technology, Inc., 1:1000), Nanog (cat. no. 4893; Cell Signaling Technology, Inc., 1:1000), ZEB1 (cat. no. sc-25388; Santa Cruz Biotechnology, Inc., 1:1000), SLUG (cat. no. sc-166476; Santa Cruz Biotechnology, Inc., 1:1000), SNAIL (cat. no. sc-10432; Santa Cruz Biotechnology, Inc., 1:1000), Twist (cat. no. sc-15393; Santa Cruz Biotechnology, Inc., 1:1000), ALDH1A1 (cat. no. ab6192; Abcam, 1:1000), ALDH1A3 (cat. no. ab129815; Abcam, 1:2000), BIRC5 (cat. no. ab76424; Abcam, 1:1000), E-Cadherin (cat. no. ab15148; Abcam, 1:1000), CD133 (cat. no. ab19898; Abcam, 1:1000), N-Cadherin (cat. no. 610920; BD Transduction, 1:1000), Vimentin (cat. no. MA5-14564; Invitrogen, 1:1000) and IGFBP-3 (cat. no. AF675; R&D SYSTEMS, 1:1000) were used for Western blot analysis and Immunocytochemistry assays.

### Small interfering RNA (siRNA) mediated knockdown of BIRC5

A549 and U87 cells were transfected with siRNA targeting BIRC5 (GAGGUUCCAAUGGCAGGUU, AACCUGCCAUUGGAACCUC; Bioneer Corporation). 10 pmol siRNAs were transfected using Lipofectamine RNAi MAX reagent (cat. no. 13-778-150; Invitrogen; Thermo Fisher Scientific, Inc.). Stealth RNAi Negative Control Medium GC (cat. no. 12935-300; Invitrogen; Thermo Fisher Scientific, Inc.) was used as the negative control. Cells were incubated at 37°C for 72 h after transfection.

### Gene expression analysis

#### RNA isolation

A-549 and NCI-H460 cell lines were treated with 1ml of TRI REAGENT (cat. no. TR118; Molecular Research Center, Inc., Cincinnati, OH 45212) for 5 minutes in the room temperature, and 200μl of Chloroform (cat. no. 28560-0350; JUNSEI, Tokyo, Japan) was added. After inverting the samples for a few times, these samples were incubated in the room temperature for 2 minutes. The samples were centrifuged (12000xg, 4°C, 15 mins) to be phased into RNAs, DNAs, and Proteins. 500μl of RNAs were separated from the sample and 500μl of Isopropanol (cat. no. 1.09634.1011; Merck KGaA, Darmstadt, Germany) were added to extract/precipitate the RNAs. After incubated in the room temperature for 5 minutes, the samples were centrifuged (1200xg, 4°C, 8 mins). Isopropanol from the samples were aspirated and the isolated RNAs were washed by using 1ml of 75% Ethyl alcohol (cat. no. 000E0690; SAMCHUN PURE CHEMICAL CO., Gyeonggi-do, Korea). 75% Ethyl alcohol were aspirated from the samples after centrifuged (7500xg, 4°C, 5mins). The isolated RNAs were dried in the room temperature for 10-15 minutes and 30-50 μl of DEPC water were added to the dried RNAs. These RNA samples were incubated in 60°C heat plate for 10-15 minutes. After heating, these samples were cooled by ice.

#### Agarose gel electrophoresis

Spectrophotometer ASP-2680 (ACTGene; Piscataway, USA) was used to measure the quantity of the extracted RNA samples. 1μg of each RNAs were added into Maxime RT Premix (Random primer) (cat. no. 25082; LiliF Diagnostics, Gyeonggi-do, Korea) to make cDNA. DEPC water were used to fulfill the total volume as 20μl. For RT-qPCR, ALDH1A1 (F : TTAGCAGGCTGCATCAAAAC, R : GCACTGGTCCAAAAATCTCC, 56°C, 34 Cycle), ALDH1A3 (F : ACCTGGAGGGCTGTATTAGA, R : GGTTGAAGAACACTCCCTGA, 57.5°C, 34 Cycle), CD133 (F : CATGGCCCATCGCACT, R : TCTCAAAGTATCTGG, 55°C, 34 Cycle), BIRC5 (F : AGAACTGGCCCTTCTTGGAGG, R : CTTTTTATGTTCCTCTATGGGGTC, 56°C, 34 Cycle), GAPDH (F : AGTCAACGGATTTGGTCGTA, R: GTCATGAGTCCTTCCACGAT, 56°C, 34 Cycle), and IGFBP-3 (F : CAGAACTTCTCCTCCGAGTCC, R : CCACACACCAGCAGAAACC, 56°C, 34 Cycle) primers were used. 18μl of DEPC water and 2μl of each primers were added in the Maxime^™^ PCR PreMix (i-MAX II for 20μl). We used T100^™^Thermal Cycler (Bio-Rad Laboratories, Inc., Hercules, CA, USA) for RT-qPCR. After the whloe steps, 0.5X TAE buffer (cat. no. IBSD-BT002; iNtRON Biotechnology, Gyeonggi-do, Korea) and Certified^™^ Molecular Biology Agarose (cat. no. 161-3102; Bio-Rad Laboratories, Inc., Hercules, CA, USA) were mixed to make 1% gel. 20μl of samples were loaded on the gels for 10-20 minutes. The loaded gels were put on the Desktop Gel Imaging System (ETX-20.M; EEC Biotech Co., Ltd.) to be pictured.

#### Western Blot Analysis

Cells were lysed in RIPA Lysis Buffer, 10X (20-188, Millipore, DARMstadt, Germany) containing phosphatase inhibitor cocktail tablets (04906837001, Roche, Basel, Swiss) and protease inhibitor cocktail tablets (11836153001, Roche, Basel, Swiss). Protein concentration was measured by Protein Assay Dye Reagent Concentrate (cat. #5000006, BIO-RAD, Hercules, CA, USA). For a Western blot analysis, equal amounts of protein were separated on 8-15% sodium dodecyl sulfate (SDS)-polyacrylmide gels and separated protein was transferred to Amersham ^™^ Protran ^™^ 0.2μm NC (10600001, Amersham^™^; Cytiva, Pittsburgh, PA, USA). After blocking transferred membrane in room temperature by using phosphate-buffered saline (PBS) buffer containing non-fat milk (10%) and Tween 20 (0.1%) for 1 hour, membranes were treated with specific antibodies overnight in a cold chamber. After washing with Tris-buffered saline (cat. no. A0027, BIO BASIC, Markham ON, Canada) membranes were treated with an HRP-linked secondary antibody (Anti-rabbit IgG cat. no. 7074S or Anti-mouse IgG cat. no. 7076S; Cell Signaling Technology, Inc) for 2 hours at room temperature, and visualized by WESTERN BLOTTING LUMINOL REAGENT (cat. no. sc-2048, Santa Cruz Biotechnology, Inc.).

#### Sphere formation assay

For sphere formation of cancer cells, conditioned media was used. Conditioned media refers to stem cell-acceptable DMEM (DMEM-F12; cat. no. 11320-033; Invitrogen; Thermo Fisher Scientific, Inc.) supplemented with basic fibroblast growth factor (bFGF; 20 ng/ml; cat. no. 13256-029; Invitrogen; Thermo Fisher Scientific, Inc.), epidermal growth factor (20 ng/ml; cat. no E9644; Sigma-Aldrich; Merck KGaA), and B27 Serum-Free Supplement (cat. no. 17504-044; Invitrogen; Thermo Fisher Scientific, Inc.).

#### Single cell assay

Single cell experiments were set up with floating cells in an ultra-low adhesion 96-well plate (cat. no. 3474; Corning, Inc.) with one or two cells distributed into each well. The cells were cultured in a humidified incubator with 5%CO_2_ at 37°C. The following day, wells with single cells in them were selected visually under a light microscope (magnification, x400), and after 10-14 days, spheres were quantified based on absolute count as well as diameter, and photographed using an inverted phase contrast microscope.

#### Limited dilution assay

In a limited dilution assay, cells were plated in 200 μl spheroid formation assay medium in ultra-low adhesion 96-well plates. A total of 1, 10, 50, 100 or 200 cells/well were plated, with 48 wells for each starting density of cells. Oncospheres were analyzed using a light microscope (magnification, x400) after 10-14 days of incubation. A well with at least one spheroid with a diameter ≥100 μm was defined as a positive well, and the number of positive wells was counted.

#### Colony-forming assay and irradiation

Cells were seeded in 35-mm culture dishes at a density of 1 × 10^3^ cells per plate and attached overnight. The next day cells were exposed to a 3-Gy dose of γ-radiation. After 10–14 days, cells were stained with 0.5% crystal violet and colonies (defined as groups of ≥50 cells) were counted. Clonogenic survival was expressed as a percentage relative to the non-irradiated controls.

#### Invasion and migration assay

An invasion assay was performed using a Matrigel-coated invasive chamber (8-μm pore; BD Biosciences) according to the manufacturer’s instructions. A migration assay was carried out using an uncoated chamber. The lower culture chamber of the 24-transwell plate (Cell Biolabs) was filled with 500 μl of a transfer medium consisting of RPMI 1640 and 10% FBS. Cells were seeded in the upper chamber at a density of 1 × 10 ^5^ cells in a 200 μl serum-free medium/well and incubated for 48 hours at 37 ° C in a humidified atmosphere of 5% CO_2_. Cotton swab was used to wipe out the non-mobile cells in the upper chamber. On the other side of the upper chamber, the moved cells were stained with crystal violet and the cells were counted with an optical microscope.

#### Wound healing assay

Cells were plated in a 60 mm culture dish and grown to 80% confluence. A wound was created by scraping the monolayer of cells with a 200 μl pipette tip in the middle of the dish. Floating cells were removed by washing with PBS and fresh medium containing 10% FBS was added. The doubling time of the A549 cells was 22 h. Cells were incubated at 37°C for 24 h, and imaged using phase-contrast microscopy (magnification, x400). The distance between the edges of the wounds shown in the image was measured randomly at three or more places and the mean of the three measurements was obtained.

#### Immunocytochemistry

Cells were grown onto glass coverlips in 35mm plates and fixed with 4%paraformaldehyde(cat. no. P2031; Biosesang) for 30mins at room temperature. After cell fixation, cells were incubated with antibodies in a solution of Tris-buffered saline (cat. no. A0027, BIO BASIC, Markham ON, Canada) at 4°C for overnight. The antibodies used were: BIRC5(cat. no. ab76424; Abcam), ALDH1A1(cat. no. ab6192; Abcam), ALDH1A3(cat. no. ab129815; Abcam), CD133(cat. no. ab19898; Abcam), CD44(cat. no. 3570; Cell Signaling Technology, Inc.), E-Cadherin(cat. no. ab15148; Abcam), N-Cadherin (cat. no. 610920; BD Transduction), Vimentin(cat. no. MA5-14564; Invitrogen). Staining was visualized using Alexa Fluor 488–conjugated anti-rabbit IgG antibody(Invitrogen). Nuclei were counterstained using 4,6-diamidino-2-phenylindole(DAPI; Sigma-Aldrich). Stained cells were analyzed using a Zeiss LSM510 Meta microscope (Carl Zeiss Micro Imaging GmbH, Göttingen, Germany).

#### Kaplan-Meier plotter

Using the published genetic information system, Kaplan-Meier survival values were obtained (kmplot.com/analysis). This was based on the results of a mRNA gene chip analysis using tissues from lung cancer patients. The gene symbol used was BIRC5. All conditions were set as total lung cancer patients. Significant results obtained using a Kaplan-Meier survival analysis are expressed as log-rank *p*-values for the significance of differences in survival between groups.

#### Cytokine array

Human XL Cytokine Array Kit(cat. no. ARY022B; R&D SYSTEMS) was used to evaluate the secreted factors regulated by BIRC5 in lung cancer stem cells. A549 cell was transfected with siRNA targeting BIRC5. All reagents were brought to room temperature before used. 2ml of blocking buffer were put into each well of the 4-Well Multi-dish and incubated for 1 hour on a rocking platform shaker. While the arrays were block, the samples were prepared by diluting the desired quantity to a final volume of 1.5ml with blocking buffer. blocking buffer were aspirated from the wells of the wells of the 4-Well Multi-dish after 1 hour, and the prepared samples were added. After incubated for overnight at 2-8 °C on a rocking platform shaker, each membrane were carefully removed and placed into plastic containers with 20mL of 1xWash Buffer. For 10 minutes, each membrane were washed with 1xWash Buffer on a rocking platform shaker for 3 times. 30μL of Detection Antibody Cocktail were added to 1.5ml blocking buffer and 1.5ml of diluted Detection Antibody Cocktail were pipetted into the 4-Well Multi-dish. After 1 hour incubation on a rocking platform shaker, each membrane were washed with 1xWash Buffer for 3 times. Each membrane were removed from the container and visualized by Chemi Reagent Mix.

#### Statistics and reproducibility

All experiments were performed with three or more biological replicates. Statistical analyses were performed using PRISM version 7.0 (Graph Pad Software, San Diego, CA, USA). Experimental data are presented as means ± standard deviation (s.d.) of five or more independent experiments. Each exact n value is indicated in the corresponding figure legend. Comparisons were performed using two-tailed paired Student’s t-tests. Not significant (ns): P > 0.05; significant.

## Results

### BIRC5 highly expressed in malignant lung cancer

In a previous study, a list of genes overexpressed in ALDH1^+^ cells, known as one of the marker proteins of lung cancer stem cells, was obtained (23). Commonly expressed genes were found by comparing the obtained gene group with the gene group obtained from GSC, and BIRC5 was among them. The genetic results obtained from GSC utilized the results of papers from other groups (24). Previous studies have shown that A549 cells are more than 80% ALDH1^+^ (23). In contrast, we found that H460 cells have very low expression of ALDH1^+^ (25). Therefore, we would like to conduct an experiment with these two cells. First, the expression level of BIRC5 was confirmed in A549 cells with high ALDH1 expression. A549 cells are known to have resistance to irradiation, whereas H460 cells are known to be sensitive to irradiation (23). Differences in gene expression of the LCSC markers ALDH1A1, ALDH1A3, and CD133 and BIRC5 were compared in A549 and H460 cells (Fig 1A). From the results, A549 cells showed higher expression than H460. Fig 1B confirms the difference in protein expression, and the results show the same pattern as Fig 1A. Fig 1C visually confirms the expression patterns of the cancer stem cell marker protein and the BIRC5 gene using ICC (immunocytochemistry). ICC results were identical to those using Western blotting (WB). Fig 1D shows that patients with high BIRC5 expression have a poor prognosis based on the Kaplan Meier survival curves.

**Fig 1.**
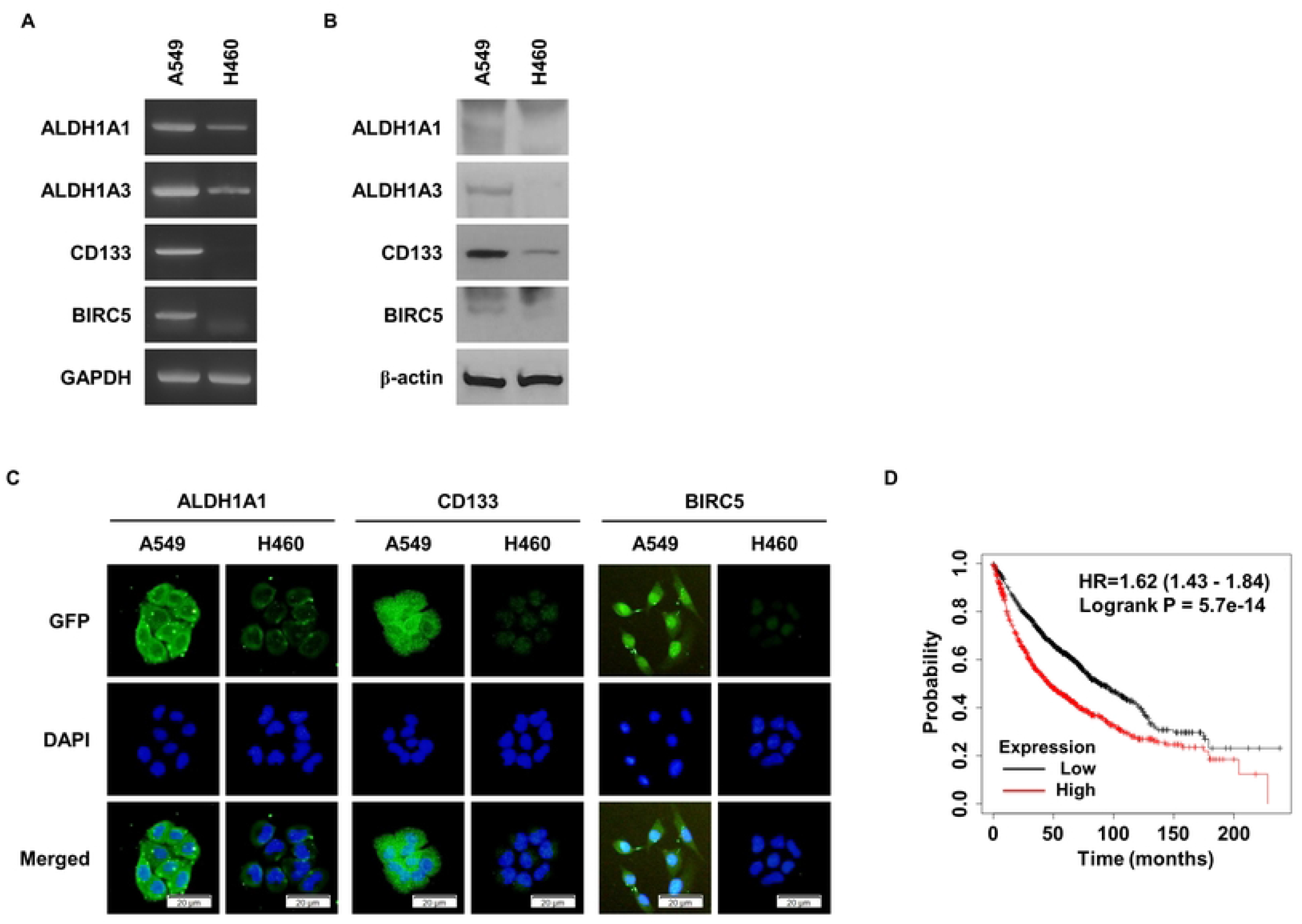
BIRC5 is expressed in A549 cells with high ALDH1 expression. (A) Comparison of CSC marker gene expression and BIRC5 gene expression in A549 and H460 cells. (B) Comparison of CSC marker protein expression and BIRC5 protein expression. (C) Comparison of CSC marker protein expression and BIRC5 expression using ICC. (D) Results of Kaplan-Meier survival analysis according to BIRC5 gene expression level in lung cancer.

### Characteristics of LSCS regulated by BIRC5

When cancer cell lines are treated with conditioned media (CM), the attached cells float and form spheres after a certain period of time. Sphere formation ability is one of the major characteristics of cancer stem cells, and it was confirmed that BIRC5 is involved in this ability (Fig 2A). Cancer stem cells have self-renewal ability and can form a cluster from a single cell (Fig 2B). A single cell assay was conducted to confirm whether BIRC5 is involved in self-renewal ability, and it was found that the ability was significantly reduced in the group in which BIRC5 expression was suppressed. As presented in Fig 2C, the results of the single cell assay were verified by conducting a limited dilution assay. It was confirmed that BIRC5 is involved in major features of CSC, and the expression of CD44 and ALDH1, which are LCSC marker proteins, was confirmed. As a result, the expression of LCSC marker proteins decreased in the group in which the expression of BIRC5 was suppressed using si-RNA. Fig 2E confirms the expression of SOX2, Oct-4, and Nanog, which are known as cancer stem cell regulatory proteins, and they were regulated by BIRC5. Expression of the CSC marker protein was visually confirmed using ICC (Fig 2F). As a result, the same pattern as seen in Fig 2d was observed. A549 cells with high ALDH1 expression are known to have relatively high radioresistance. Therefore, after suppressing the expression of BIRC5, radiation was irradiated (Fig 2G). It was found that colony formation was reduced only by inhibition of BIRC5, and colony formation was significantly reduced after irradiation in the group in which BIRC5 gene was inhibited.

**Fig 2.**
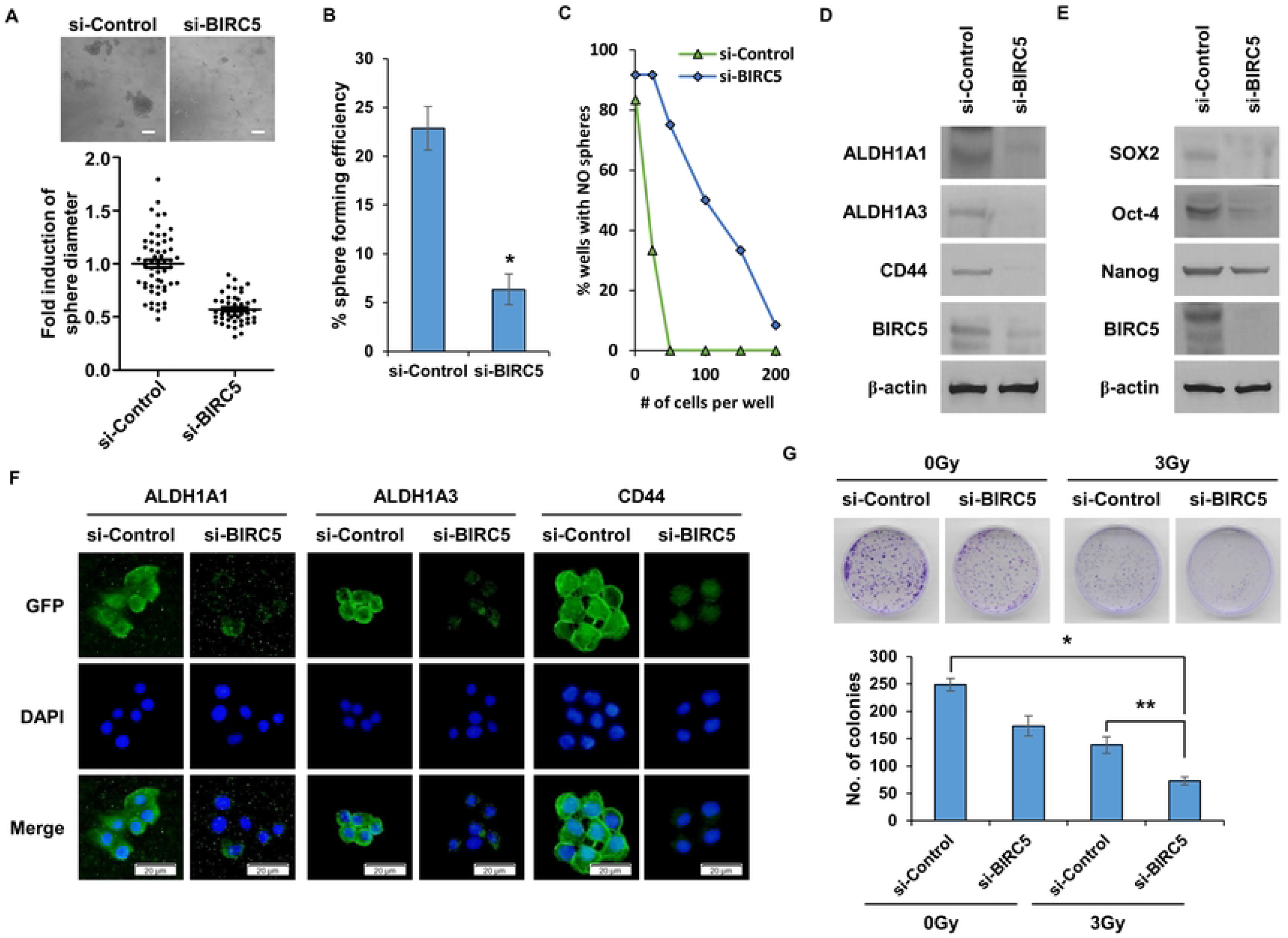
Characteristics of LCSCs regulated by BIRC5 expression. (A) Sphere formation proceeds after si-BIRC5 treatment in A549 cells. Observed seven days after CM treatment. (B) After inhibiting the expression of BIRC5 in A549 cells using si-RNA, a single cell analysis was performed. Incubated for 10 days using CM in a 96-well plate. (C) Limited dilution assay results of A549 cells treated with si-BIRC5. Using a 96-well plate, 1, 25, 50, 100, 150, and 200 cells were put into one well and incubated. (D) Expression of CSC marker proteins in A549 cells in which BIRC5 expression was suppressed by si-RNA treatment. (E) Expression of CSC regulator proteins in A549 cells in which BIRC5 expression was suppressed. (F) CSC marker protein expression using ICC. (G) Colony forming assay to determine whether BIRC5 is involved in radioresistance. 3Gy radiation was irradiated. Data are presented as the mean ± standard deviation of three repeats. Scale bar, 50μm. *p<0.001, **p<0.005.

### BIRC5 regulates cell motility and EMT phenomenon

Cancer stem cells are known to be closely related to the EMT phenomenon (26–28). Therefore, the regulation of the EMT phenomenon by BIRC5 was confirmed. First, after suppressing the BIRC5 gene, the expression of E-cadherin, N-cadherin, and Vimentin, which are marker proteins for EMT, was evaluated using WB (Fig 3A). The expression of E-cadherin, an epithelial marker protein, increased in the BIRC5-inhibited group, while the expression of mesenchymal marker proteins N-cadherin and Vimentin decreased. Fig 3B shows whether the expression of Snail, Slug, Twist, and Zeb1, which are EMT regulatory proteins, is related to BIRC5. It was confirmed that in the group in which the expression of BIRC5 was decreased, all proteins regulating the EMT phenomenon were decreased. As a result of evaluating the expression of the EMT phenomenon marker protein using ICC, the same result as in Fig 3A was obtained (Fig 3C). The most notable feature of the EMT phenomenon is the enhancement of cell migration ability (29–31). A migration and invasion assay was performed using a Boyden chamber to confirm whether BIRC5 affects the migration ability of cells (Fig 3D). In the group in which the BIRC5 gene was suppressed, both cell migration and invasion abilities were reduced. As determined by conducting a wound healing assay to confirm the cell’s migration ability, the cell’s migration ability was significantly reduced, consistent with the result obtained using the Boyden chamber (Fig 3E).

**Fig 3.**
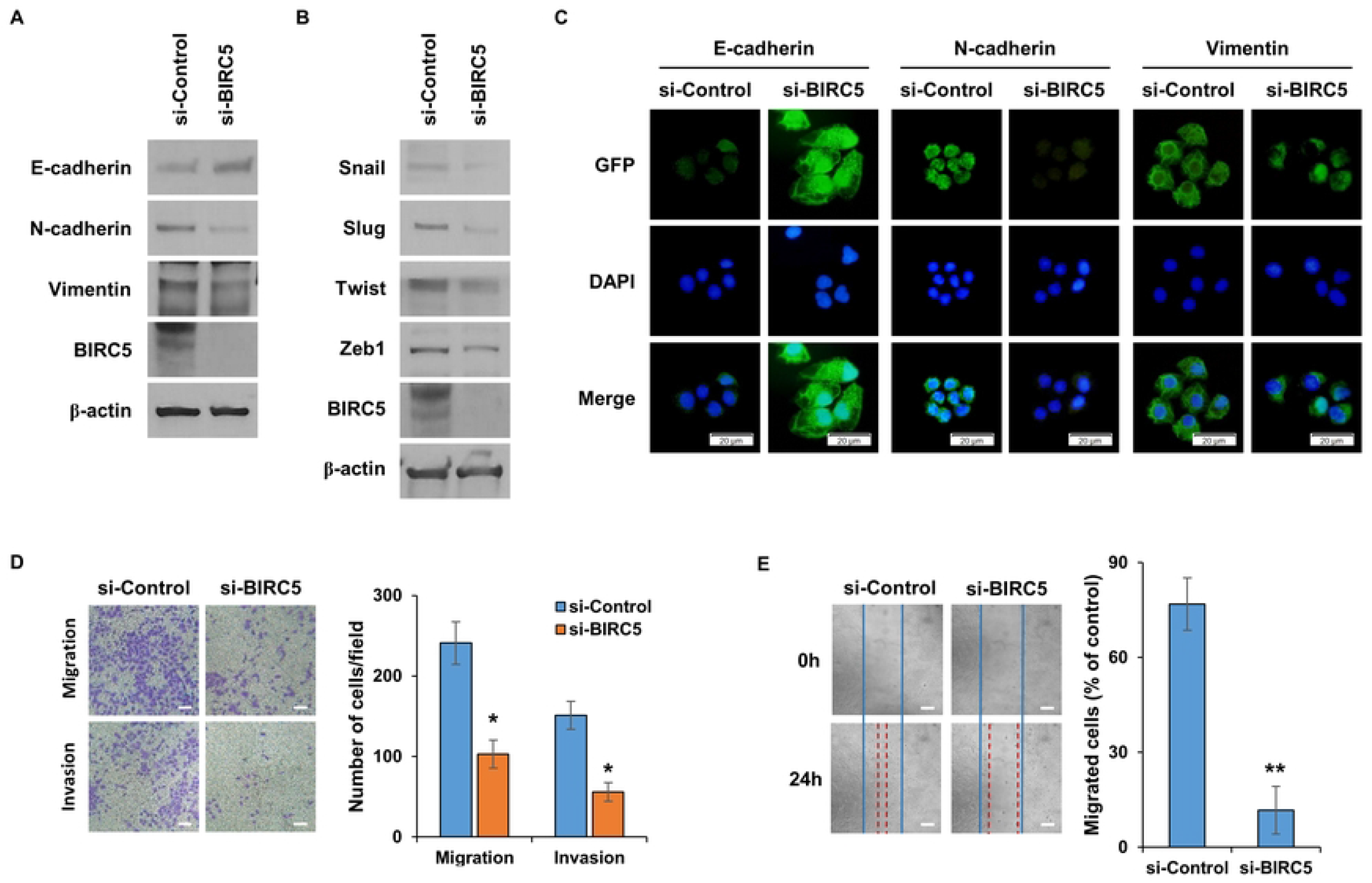
BIRC5 is involved in EMT in LCSCs. (A) Expression patterns of EMT marker proteins according to inhibition of BIRC5 expression. (B) Changes in the expression patterns of EMT regulatory proteins according to the inhibition of BIRC5 expression by si-RNA treatment. (C) Differences in EMT marker protein expression according to the expression level of BIRC5 gene using ICC. (D) Cell migration and invasion ability was confirmed after si-BIRC5 treatment. Results confirmed 48 hours after inoculation of A549 cells into a Boyden chamber. (E) Wound healing assay to evaluate the difference in cell motility according to the expression of BIRC5. Observed 24 hours after wounding of A549 cells. Data are presented as the mean ± standard deviation of three repeats. Scale bar, 50μm. *p<0.005, **p<0.0001.

### Regulate malignant traits of cancer through secreted factor IGFBP-3

Cancer stem cells have signal transduction mechanisms by exogenous secreted factors as well as intracellular signal transduction mechanisms (32, 33). Therefore, as a result of evaluating the secreted factors regulated by BIRC5, the expression of IGFBP-3 was significantly reduced (Fig 4A). The gene expression level of IGFBP-3 according to the expression level of BIRC5 was confirmed using PCR (Fig 4B). When the gene expression of BIRC5 is suppressed, the gene expression of IGFBP-3 is also suppressed. After suppressing the function of IGFBP-3 using an IGFBP-3 neutralizing antibody, expression of BIRC5 was confirmed (Fig 4C). The results revealed that the expression of BIRC5 was suppressed as IGFBP-3 decreased. This confirmed that BIRC5 and IGFBP-3 form a single ring. Fig 4d shows that the expression of ALDH1 and CD44, which are CSC marker proteins, decreased after the use of the IGFBP-3 neutralizing antibody. In addition, as a result of checking the expression level of the EMT marker protein after inhibiting the function of IGFBP-3 with a neutralizing antibody, it was confirmed that the EMT phenomenon was regulated by IGFPB-3 (Fig 4E).

**Fig 4.**
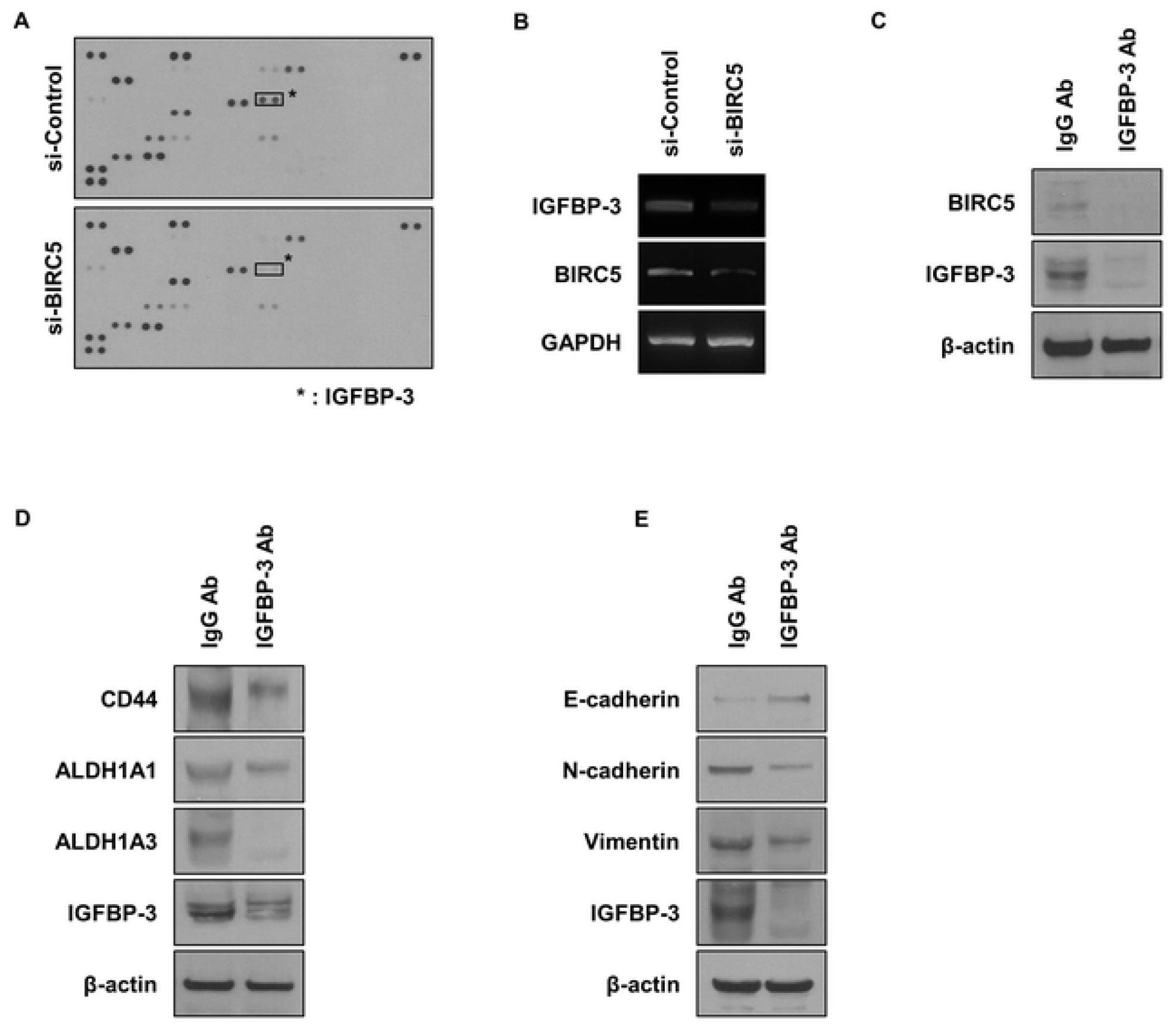
BIRC5 regulates the secretion of IGFBP-3. (A) Analysis of differences in secreted factors according to the expression of BIRC5. (B) Confirmation of the gene expression level of IGFBP-3 when the expression of BIRC5 was inhibited using si-RNA. (C) Confirmation of BIRC5 expression when IGFBP-3 was inhibited by treatment with a neutralizing antibody. (D) Confirmation of difference in expression of CSC marker protein after treatment with neutralizing antibody. (E) After suppressing IGFBO-3, the expression level of EMT marker protein was determined.

### GSC and EMT regulated by BIRC5 in glioma

BIRC5 was highly expressed in GSCs as well as LCSCs. Therefore, the function of BIRC5 in GSC was confirmed using U87 cells, a glioma cell line. After suppressing the expression of BIRC5 in U87 cells using si-RNA, differentiation of cancer stem cells was induced by treatment with CM. In the group in which BIRC5 expression was suppressed, the expression of ALDH1 and CD133, which are CSC marker proteins, decreased (Fig 5A). Through visual confirmation using ICC, the same results as presented in Fig 5A were obtained (Fig 5B). Fig 5C measures the sphere-forming ability, which is a characteristic of CSCs. In the group with low BIRC5 expression, the ability to form spheres was found to be reduced. After inhibiting the expression of the BIRC5 gene, the expression of E-cadherin, N-cadherin, and Vimentin, which are EMT marker proteins, was assessed (Fig 5D). From the results, the expression of E-cadherin increased and the expression of N-cadherin and Vimentin decreased, similar to the LCSC results. This was consistent with the experimental results using ICC (Fig 5E). Cell migration and invasion ability, which are the main features of the EMT phenomenon, was also investigated (Fig 5F). It was found that both the migration and invasion abilities of cells were regulated by BIRC5. Finally, the expression level of IGFBP-3 regulated by BIRC5 in LCSC was confirmed in GSC (Fig 5G). It was thus confirmed that IGFBP-3 was regulated by BIRC5 in GSC as in LCSC.

**Fig 5.**
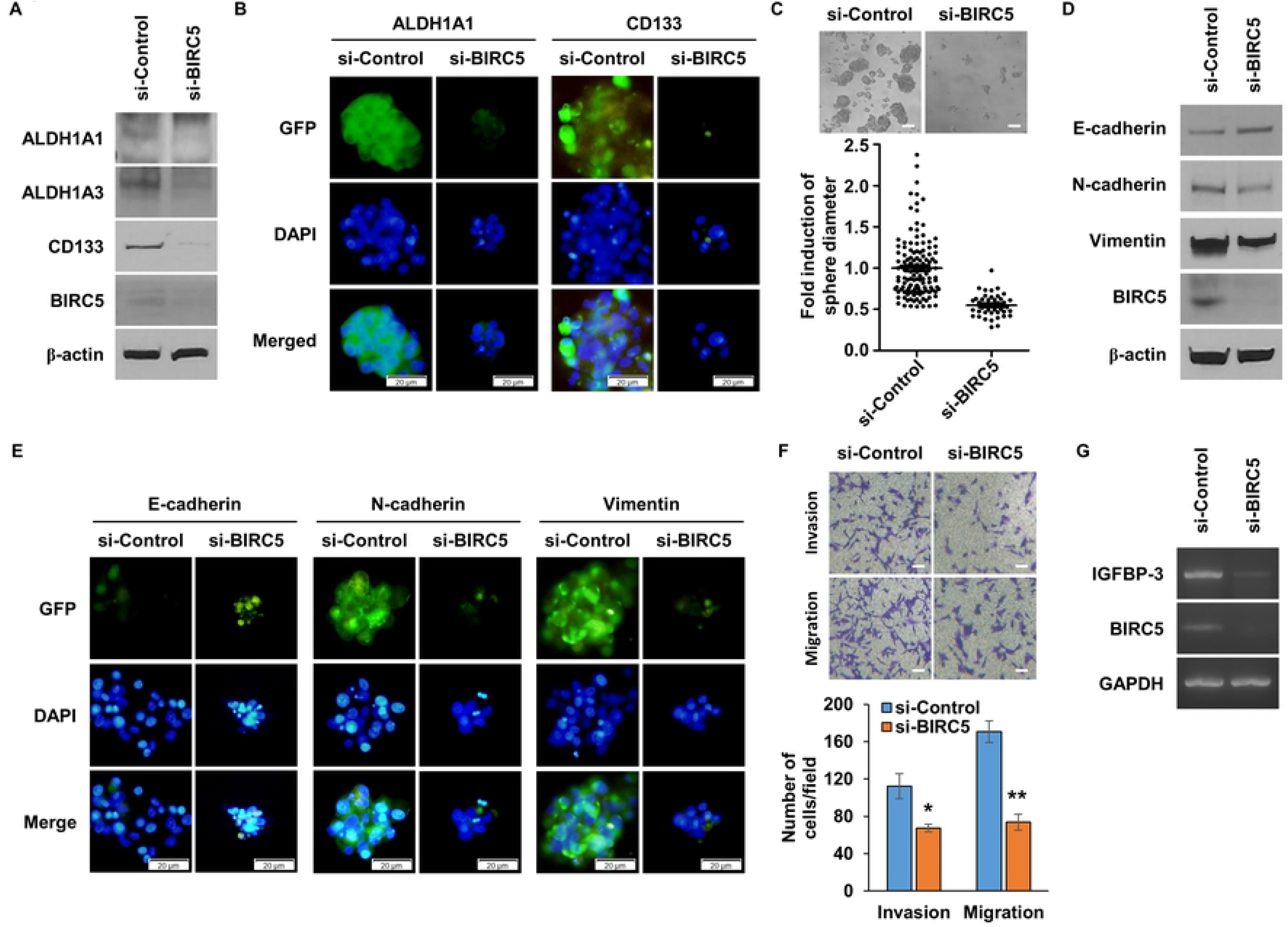
CSC and EMT regulated by expression of BIRC5 in glioma cells. (A) Expression of CSC marker protein was assessed after inhibiting BIRC5 in glioma U87 cells. (B) Visual confirmation of changes in CSC marker proteins through ICC in the cell population inhibited by si-RNA. (C) Evaluation of the degree of sphere formation in U87 cells after si-BIRC5 treatment. Observed 7 days after CM treatment. (D) A single cell analysis was performed after inhibiting the expression of BIRC5 in U87 cells. After seeding CM-treated U87 cells in a 96-well plate, one cell per well, check the results 10 days later. (E) Evaluation of changes in EMT marker proteins using si-RNA-treated U87 cells. (F) Visual confirmation of EMT marker protein changes in U87 cells in which the BIRC5 gene was suppressed through ICC. (G) An invasion and migration assay was performed after inhibiting BIRC5 by treating si-RNA. Results were obtained 48 hr after seeding the cells in the Boyden chamber. (H) Confirmation of IGFBP3 expression pattern after inhibiting the BIRC5 gene in U87 cells. Data are presented as the mean ± standard deviation of three repeats. Scale bar, 50μm. *P<0.01, **P<0.0005.

## Discussion

In this paper, it was shown that the BIRC5 gene is overexpressed in LCSC and GSC. The expression of CSC marker proteins ALDH1, CD133, and CD44 was regulated by BIRC5, and CSC regulators Sox2, Oct4, and Nanog were also regulated. In addition, sphere formation ability, self-renewal, and radiation resistance, which are known as characteristics of CSC, were all regulated by BIRC5. CSCs regulated by BIRC5 were regulated not only by LCSCs but also by GSCs. BIRC5 is known to be involved in signaling pathways including PI3K/AKT, JAK/STAT, WNT/-Catenin, and NOTCH (34). This signaling mechanism is considered the most important signaling mechanism of CSCs (35, 36). The detailed signaling mechanism of BIRC5 will be confirmed through additional studies.

IGFBP-3 was found to be regulated by BIRC5. IGFBP-3 is known to play a role in assisting IGF, a growth factor. IGFBPs consist of six families, all of which have greater binding affinity to IGF than to IGF-1R. Specifically, IGFBP-3 binds to glycoproteins in the circulatory system and slows the clearance of IGF with a very long half-life. IGFBP-3 also enhances the half-life of IGF, but due to its high binding affinity with IGF, it can inhibit cell proliferation and survival (37). IGFBP-3 enhances the adhesion of normal breast epithelial cells by binding to the matrix and activates the MAPK signaling pathway (38). It is not known what role IGFBP-3, which is involved in the MAPK signaling pathway through substrate binding, plays in breast cancer. What is currently known is that IGFBP-3 as an additive inhibits the attachment of prostate cancer cells to specific components of the extracellular matrix (39). We found that IGFBP-3, which is regulated by BIRC5, forms a reciprocal loop. When we inhibited BIRC5, not only was the expression of IGFPB-3 inhibited, but when we inhibited the signaling of IGFPB-3 by treating it with a neutralizing antibody, we found that the expression of BIRC5 was inhibited. We even found that the expression of IGFPB-3 was reduced by IGFBP-3 neutralizing antibody treatment. Therefore, we plan to conduct a large-scale study of IGFs and BIRC5 that bind to IGFBP-3 and its downstream signaling.

Our findings indicate that BIRC5 is an important factor involved in the maintenance and generation of LCSCs and GSCs. Overexpression of BIRC5 has been reported in many carcinomas, and it is thought to play an important role in CSCs of carcinomas other than LCSC and GSC. In addition, IGFBP-3, whose expression is regulated by BIRC5, is known to have two functions. Although it is involved in cell death, it is also known to be involved in cell growth and cancer malignancy. The perspective presented in this paper is that IGFBP-3 is involved in cancer malignancy. Although further studies are needed, BIRC5 and IGFBP-3 are attractive new cancer treatment targets.

## Acknowledgements

This work was supported by a National Research Foundation of Korea (NRF) grant funded by the Korean government (MSIT) (NRF-2019R1C1C1009617)

